# NDC80 clustering modulates microtubule dynamics under force

**DOI:** 10.1101/198291

**Authors:** Vladimir A. Volkov, Pim J. Huis in't Veld, Marileen Dogterom, Andrea Musacchio

## Abstract

Multivalency, the presence of multiple interfaces for intermolecular interactions, underlies many biological phenomena, including receptor clustering and cytosolic condensation. One of its ultimate purposes is to increase binding affinity, but systematic analyses of its role in complex biological assemblies have been rare. Presence of multiple copies of the microtubule-binding NDC80 complex is an evolutionary conserved but poorly characterized feature of kinetochores, the points of attachment of chromosomes to spindle microtubules. To address its significance, we engineered modules allowing incremental addition of NDC80 complexes. The modules’ residence time on microtubules increased exponentially with the number of NDC80 complexes. While modules containing a single NDC80 complex were unable to track depolymerizing microtubules, modules with two or more complexes tracked depolymerizing microtubules and stiffened the connection with microtubules under force. Cargo-conjugated modules of divalent or trivalent NDC80 stalled and rescued microtubule depolymerization in a force-dependent manner. Thus, multivalent microtubule binding through NDC80 clustering is crucial for force-induced modulation of kinetochore-microtubule attachments.

---

Many macromolecular interactions engage binding partners with multiple subunits (oligomers) or interaction interfaces. High binding valency (multivalency) allows simultaneous interactions that result in a multiplying effect on binding affinity, as shown by the classic example of immunoglobulins, which recognize foreign antigens in the immune response.

Multivalent interactions may occur in solution or on defined interfaces, such as cellular membranes or chromosomes. Kinetochores are multiprotein assemblies built on the centromeres of chromosomes. Their ability to bind microtubules, dynamic polymers that alternate between phases of growth and shrinkage, is crucial for the segregation of the replicated chromosomes (sister chromatids) to the daughter cells during cell division. The 4-subunit NDC80 complex (NDC80, for nuclear division cycle 80 complex) provides the crucial link between kinetochores and microtubules. Quantitative fluorescent microscopy and bottom-up reconstitutions indicated that kinetochores contain multiple NDC80 complexes, with recent estimates converging on 8 complexes per microtubule-binding site (Musacchio and Desai, 2017). In humans, a single kinetochore contains a molecular lawn of approximately 200 NDC80 complexes that bind 15-20 microtubules (Suzuki et al., 2015; Wendell et al., 1993). With hundreds of different proteins assembled on a chromosome’s centromeric region, kinetochores are the epitome of a multivalent proteinaceous platform, but how the stoichiometry and modular organization of NDC80 contribute to the coupling of kinetochores to microtubules remains unclear.

To address this question, we reconstituted kinetochore modules containing one, two, three, or four copies of human NDC80. Traptavidin (T), a streptavidin variant with tenfold slower biotin dissociation, and a biotin-binding deficient streptavidin variant tagged with a SpyCatcher module (S) were folded into TS tetramers with stoichiometries varying from T_4_S_0_ to T_0_S_4_ (Chivers et al., 2010; Fairhead et al., 2014; Zakeri et al., 2012) (Figure S1). Purified TS tetramers were then covalently coupled to NDC80 complexes that contain SPC24^SpyTag^ and fluorescently labeled SPC25. Assemblies containing one, two, three, or four NDC80 complexes were subsequently separated by size-exclusion chromatography chromatography and analyzed by SDS-PAGE (Figure 1A and Figure S1). Inspection of the purified TS-NDC80 modules by electron microscopy after glycerol spraying and low-angle metal shadowing confirmed the integration of elongated NDC80 tethers on a central TS density (Figure 1B-C). The flexible orientation of the NDC80 complexes is highly reminiscent of NDC80 clustered on their kinetochore receptors CENP-C and CENP-T (Huis in’t Veld et al., 2016). We thus reconstituted a proxy of the outer kinetochore with precise control over the number of incorporated NDC80 complexes.

**Figure 1.**
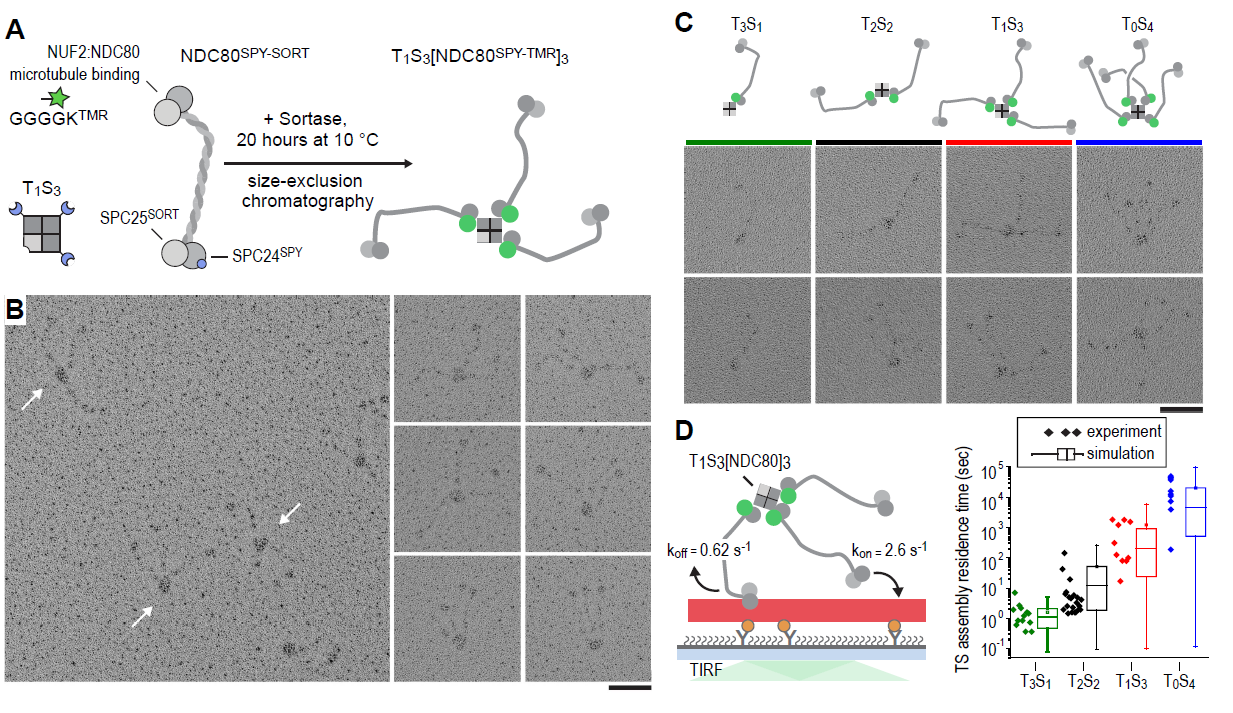
Incremental addition of NDC80 results in hyperstable microtubule binding. **a**) NDC80^SPY-SORT^ was fluorescently labelled and covalently bound to TS assemblies. The cartoon shows the formation of T_1_S_3_[NDC80]_3_ assemblies. Size-exclusion chromatography and SDS-PAGE analysis are shown in Figure S1. **b**) T_1_S_3_[NDC80]_3_ assemblies were analysed by electron microscopy after low-angle rotary shadowing. Three flexible NDC80 complexes of approximately 60 nm originate from central T_1_S_3_ densities (white arrows in the field of view) Scale bar 50 nm. **c**) Side-by-side comparison of NDC80 coupled to T_3_S_1_, T_2_S_2_, T_1_S_3_, and T_0_S_4_. Cartoons represent the approximate orientation of assemblies in the upper row of micrographs. Scale bar 50 nm. **d**) Residence time of quantized NDC80 assemblies on taxol-stabilized microtubules as determined experimentally (dots) and as predicted by a series of 1000 simulations (box and whiskers plot; box: 25-75%, horizontal line: median, whiskers: 5-95%). NDC80 complexes of a microtubule-bound TS-NDC80 assembly attach to and detach from microtubules with rates of *k*_*on*_ and *k*_*off*_, respectively. The residence time of an oligomer is defined as the time between the association of its first NDC80 tether and the detachment of all NDC80 tethers.

To test at a single-molecule level how assemblies with a precisely defined NDC80 stoichiometry interact with microtubules, we measured the residence time of TS-NDC80 modules on taxol-stabilized microtubules using total internal reflection fluorescence (TIRF) microscopy (Figure 1D and Figure S2). The residence time of these assemblies on microtubules increased more than tenfold for every additional NDC80 complex (Figure 1D). To simulate the binding of TS-NDC80 assemblies to microtubules *in silico*, we assumed transitions between the number of microtubule-bound NDC80 tethers based on the *k*_on_ and *k*_off_ rates of each individual NDC80. The initial landing rate of an assembly on a microtubule was ignored so that each simulation started with a single NDC80 bound to a microtubule and stopped after detachment of all NDC80s. Using the reciprocal of the residence time of T_3_S_1_[NDC80]_1_ as *k*_*off*_, stochastic simulations of the residence times of mono-, di-, tri-, and tetravalent assemblies were used to determine *k*_*on*_ (Figure 1D). A fit to the data resulted in a *k*_*on*_ of 2.6 s^-1^. Assuming a TS-NDC80 assembly as a sphere, one NDC80 inside the sphere has a local concentration of 1.7 µM. This predicts a concentration-dependent association rate of 1.5 µM^-1^s^-1^, a value that falls well within the published range of *k*_*on*_ for free NDC80 binding to microtubules (Powers et al., 2009; Tien et al., 2010; Zaytsev et al., 2015). Taken together, this demonstrates that all NDC80 complexes in TS-NDC80 modules can interact with one microtubule and illustrates how the clustering of NDC80 stabilizes microtubule binding.

We next used dynamic microtubules to test how assemblies with multiple NDC80 tethers interact with microtubule ends. For this purpose, TS-NDC80 assemblies, tubulin, and GTP were added to GMP-CPP stabilized microtubule seeds attached to passivated coverslips (Figure 2A). Using TIRF microscopy, fluorescently labelled TS-NDC80 was imaged on dynamic microtubule extensions (Figure 2B and Video 1). Consistent with previous findings (Powers et al., 2009; Schmidt et al., 2012), monomeric NDC80 dissociated from depolymerizing microtubules. Di-, tri-, as well as tetravalent assemblies tracked shortening microtubule tips and each additional NDC80 increased the fraction of tip-tracking assemblies (Figure 2C and Figure S2). Free microtubules in the same flow chamber shortened faster than the ones with TS-NDC80 at the tip (Figure 2D). The ability to impede microtubule depolymerization suggests that tip-tracking TSNDC80 modules form a direct connection to the shortening microtubule ends.

**Figure 2.**
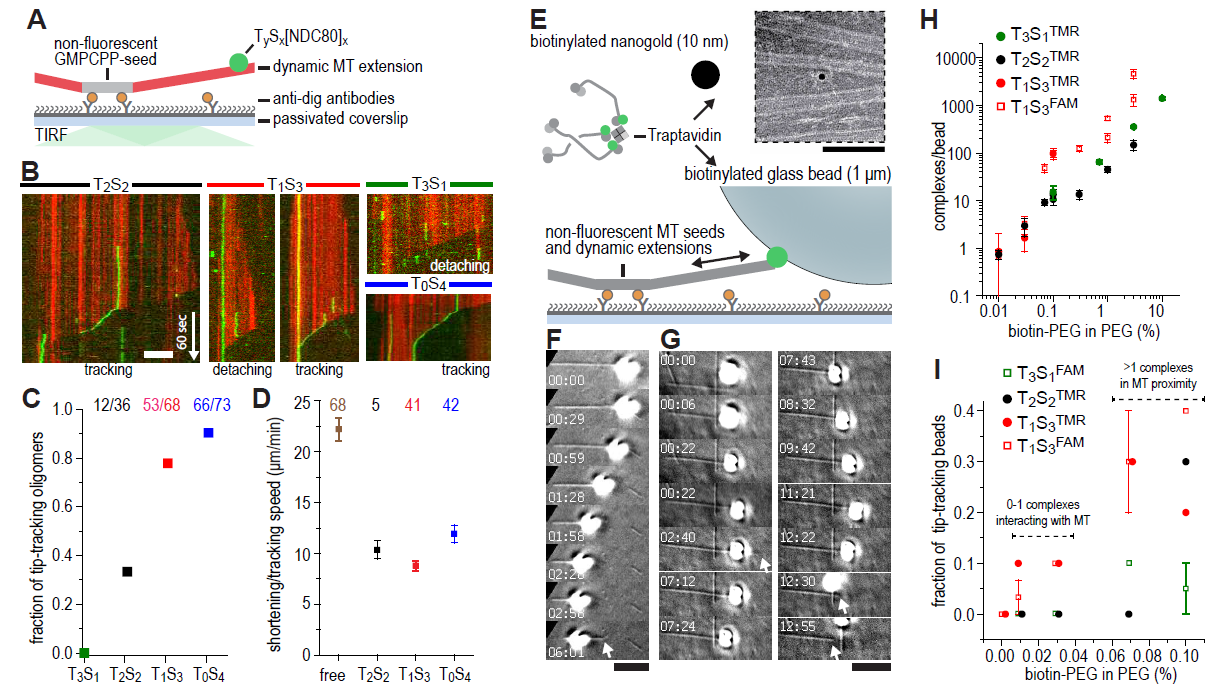
Trivalent TS-NDC80 efficiently tracks depolymerizing microtubules and transports cargo. **a**) Schematic representation of the experimental setup. **b**) Kymographs showing NDC80 (green) assembled on T_2_S_2_, T_1_S_3_, or T_0_S_4_ tracking a depolymerizing microtubule (red). An example of a T_1_S_3_[NDC80]_3_ complex that detached from the tip of shortening microtubule is also included. Scale bar 5 µm. See Figure S2 and Video 1. **c**) The fraction of NDC80 assemblies that track depolymerizing microtubules. **d**) Comparison between microtubule depolymerization in the presence and absence of TS-NDC80 following the shortening tips. Data are shown as mean ± SEM. **e**) Biotinylated glass beads or nanogold particles can be conjugated to traptavidin in TS-NDC80C assemblies. Nanogold particles coated with T_1_S_3_[NDC80]_3_ bound to microtubules as observed by negative-staining EM (see also Figure S3). Scale bar 100 nm. **f-g**) Examples of glass beads coated with T_1_S_3_[NDC80]_3_ tracking depolymerizing microtubules. The bead in panel g follows the growing microtubule after a rescue event until it detaches during a second depolymerization phase. White arrows indicate the dynamic microtubule tips. Scale bar 5 µm. **h**) Fluorescence-based quantification of the number of complexes on glass beads coated with increasing amounts of PLL-PEG-biotin. **i**) The fraction of beads coated with various NDC80 assemblies that track depolymerizing microtubules as a function of the amount of biotin-PEG added to the beads (see also Figure S4).

We next set out to test if TS-NDC80 can couple forces generated by the depolymerizing protofilaments to the movement of cargo. Biotinylated nanogold particles conjugated to trivalent TS-NDC80 modules localized on or in between microtubules, indicating that biotinylated cargo was coupled to TS-NDC80 and that cargo-bound assemblies retained their ability to bind microtubules (Figure 2E). To directly assess force coupling, we attached a biotinylated glass bead amenable to optical trapping to the biotin-binding traptavidin (T) of TS-NDC80 modules (Figure 2E and Figure S3). Binding of TS-NDC80 resulted in beads that were able to track shortening (Figure 2F) or -in rare cases-growing (Figure 2G) microtubule ends.

To control the number of TS-NDC80 assemblies on the beads, we coated beads with a mixture of poly-L-lysine-polyethylene glycol (PLL-PEG) and PLL-PEG-biotin. Using the fluorescence of NDC80C^TMR^ or NDC80C^FAM^ as a readout, we observed that the number of complexes on a bead’s surface increased linearly from <2 complexes at 0.01% PEG-biotin to several thousand complexes at 10% PEG-biotin (Figure 2H). With a diameter of 1 µm, coating of a glass bead with 0.01%-0.03% biotin-PEG predicts a single TS-NDC80 module interacting with a microtubule, whereas multiple modules are predicted to reside in the proximity of a microtubule for beads decorated with biotin-PEG concentrations above 0.07%. The presence of multiple NDC80 tethers in the proximity of a microtubule resulted in beads that followed dynamic microtubule tips for trivalent T_1_S_3_[NDC80]_3_ (13/45 cases), but sometimes also for monovalent T_3_S_1_[NDC80]_1_ (2/30 cases) (Figure 2I and Figure S4). This is consistent with the reported tip-tracking of beads that are densely decorated with monomeric NDC80 (McIntosh et al., 2008; Powers et al., 2009). Beads with the lowest biotin densities coated with monovalent T_3_S_1_[NDC80]_1_ or divalent T_2_S_2_[NDC80]_2_ were unable to follow microtubule tips (0/50 cases). The same coating densities of T_1_S_3_[NDC80]_3_ resulted in 5/70 beads following dynamic microtubule tips (Figure 2I and Figure S4). This suggests that a single trivalent T_1_S_3_[NDC80]_3_ assembly is sufficient to couple microtubule shortening to cargo transport.

We next set out to probe the force-coupling properties of TS-NDC80 assemblies with an optical trap. For this purpose, a bead coated with TS-NDC80 was positioned near the end of a dynamic microtubule, confirmed to bind the microtubule with the trap switched off, and monitored with the trap switched on while the tip of a shortening microtubule reached the bead (Figure 3A). Displacement of a bead from the centre of the trap was recorded with a quadrant photo detector (QPD). A peak in the QPD signal along the microtubule’s direction marks that the force generated by a depolymerizing microtubule is converted into the displacement of a bead coated with TS-NDC80 (Figure 3B and Video 2). After initial movement with the shortening microtubule tip, the beads stalled for an average of 1.5 ± 0.2 s (n = 91). We interpreted these stalls as the time in which the microtubule-generated force is counterbalanced by the force acting to return the bead to the centre of the trap, thus reducing the depolymerization speed to zero. Beads at the ends of stalled microtubules either detached from the microtubule and snapped back into the centre of the trap, or rescued microtubule depolymerization and gradually moved back to the centre of the trap along with the growing microtubule (Figure 3C and Video 3). In 13 out of 104 traces the bead detached from the microtubule before stalling; these traces were not analysed further.

**Figure 3.**
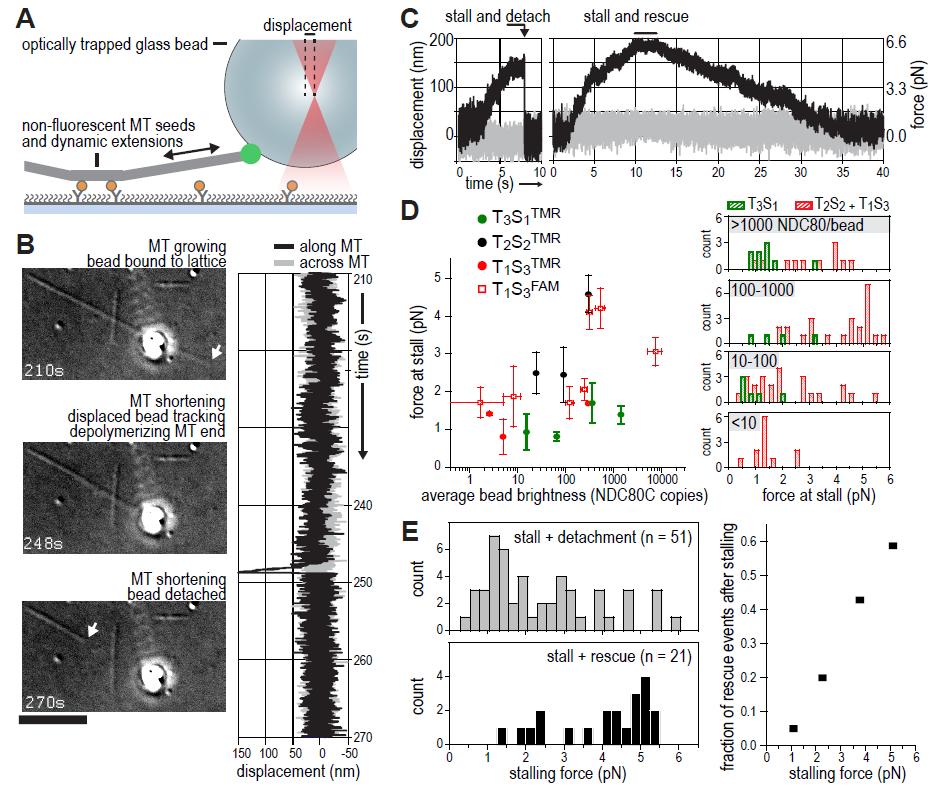
TS-NDC80 modules stall and rescue microtubule depolymerization. **a**) The displacement of an optically trapped glass bead can be used to determine the force exerted by a shortening microtubule on a bead-bound TS-NDC80 oligomer. **b**) Example of a trapped glass bead that is displaced along the microtubule axis as it holds on to a depolymerizing microtubule (248s). Arrows point to the dynamic microtubule tip before (210s) and after (270s) the force development. The graph on the right shows unfiltered QPD signal along and across the microtubule axis. **c**) Examples of unfiltered QPD signals recorded during microtubule shortening. Stalling of microtubule depolymerization by the coupled bead in the optical trap is followed by detachment of the bead (left) or a rescue of microtubule growth (right). **d**) Average forces at which differently coated beads stall shortening microtubules. Data are shown as mean ± SEM. **e**) Distribution of stalling forces that were followed by bead detachment from the microtubule (grey bars) or microtubule rescue (black bars). These distributions were used to calculate the fraction of events leading to a force-induced rescue (right).

We next tested the relation between the number of bead-bound TS-NDC80 modules and the stalling forces. Single trivalent T_1_S_3_[NDC80]_3_ stalled microtubule depolymerization at about 1.5 piconewtons (pN). Stalling forces increased to maximal values of up to 5-6 pN with 100-1000 T_1_S_3_[NDC80]_3_ modules per bead, but did not increase further with thousands of such assemblies on a single bead (Figure 3D). These beads have an estimated minimum of 4 T_1_S_3_[NDC80]_3_ modules in the proximity of a microtubule end. This organization might be optimal to couple the energy from microtubule depolymerization into cargo movement: increasing the local NDC80C concentration on the beads beyond that did not result in higher stalling forces.

Pooling of stall forces from beads coated with varying surface densities of T_1_S_3_[NDC80]_3_ and T_2_S_2_[NDC80]_2_ resulted in 51 detachment events (71%) and 21 rescue events (29%) and revealed a striking correlation between the amplitude of the stalling force and the probability of a rescue event (Figure 3E). Interestingly, even a very dense coating of monovalent T_3_S_1_[NDC80]_1_ on the beads did not generate stalling forces higher than 3 pN (Figure 3D) and never rescued microtubule shortening. Taken together, these results show that the controlled clustering of NDC80 promotes efficient coupling of microtubule-generated force as well as force-dependent rescue of microtubule shortening.

The ability of TS-NDC80 modules to slow down, stall, and rescue microtubule depolymerization suggests that they interact with the very end of the shortening microtubule. To further assess the mechanical properties of this interaction, we analysed the thermal fluctuations of trapped beads before and during contact with the tip of the microtubule. For a trapped bead in solution, these fluctuations are limited by the stiffness of the trap. For a trapped bead attached to a microtubule, these fluctuations additionally reflect the mechanical properties of the microtubule-bead linkage. During stalling, fluctuations along the microtubule were dampened compared to the signal across the microtubule and compared to fluctuations before and after the microtubule pulled on the bead (Figure 4A). This demonstrates an effective stiffening of the link between the bead and the microtubule under force. To test if pulling on microtubules and NDC80 also altered fluctuations of beads that do not interact with the depolymerizing ends, we attached beads decorated with T_1_S_3_[NDC80]_3_ assemblies either to the lattice of a microtubule away from the dynamic end, or end-on to the stabilized end of a microtubule with a GMP-CPP cap (Figure 4A). Force in these experiments was exerted by pulling on the coverslip-attached microtubule seed using the piezo stage in the same direction as a depolymerizing microtubule would pull. Stiffening of the bead-microtubule link under force was observed in all cases, but was on average two times higher during stalls produced at the interface between TS-NDC80 modules and depolymerizing microtubules (Figure 4B). Apparently, force-induced stiffening of the link provided by a cluster of NDC80 complexes is enhanced when NDC80 molecules are interacting with depolymerizing, presumably flared protofilaments, compared to when they are interacting with straight protofilaments.

**Figure 4.**
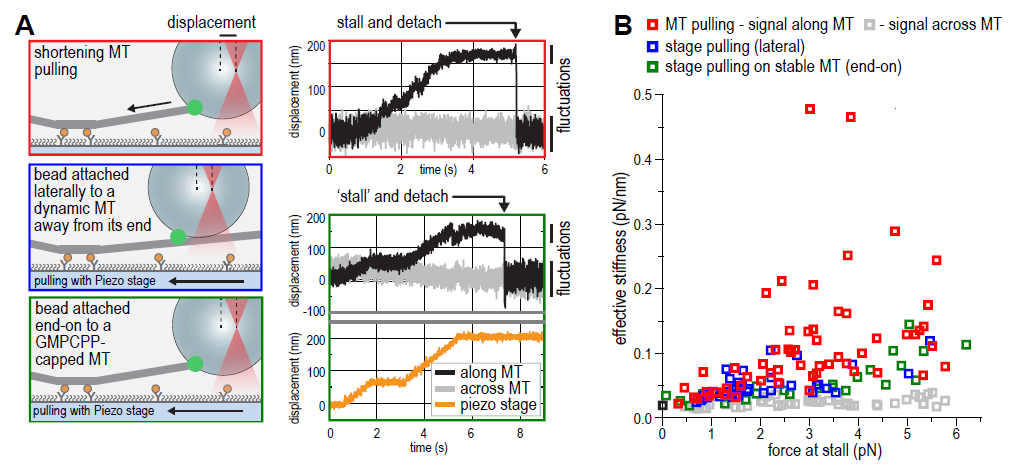
NDC80C oligomers stall microtubules through interaction with the shortening microtubule end. **a**) Experimental setup to compare forces generated by shortening microtubules (red box) with forces generating by a moving stage while a bead with T_1_S_3_[NDC80]_3_ is attached laterally to a dynamic microtubule (blue box), or end-on to a stabilized microtubule (green box). Examples of unfiltered QPD signals recorded during force generation are shown on the right. **b**) Effective stiffness of the link between the bead and the microtubule increases with force.

Kinetochore-associated fibrils that modulate the shape of depolymerizing microtubules at attached kinetochores have previously been identified *in vivo* using electron tomography, and force-induced straightening of protofilament flares has been suggested as a mechanism to convert the energy of microtubule depolymerization into cargo transport (McIntosh et al., 2008). Since suppressing the bending of protofilaments slows down microtubule shortening (Franck et al., 2007; Grishchuk et al., 2005), the impaired depolymerization rate of microtubules with plus-end tracking TS-NDC80 assemblies supports a direct binding of force-coupling NDC80 tethers to flaring protofilaments (Figure 2D). Protofilaments at growing microtubule ends are less bent than at shortening ends (McIntosh et al., 2008), therefore the force-induced protofilament straightening also explains the correlation between the magnitude of the TS-NDC80 mediated stalling force and the probability of a microtubule rescue event (Figure 3E).

Stabilization of kinetochore-microtubule attachments *in vivo* is tension-dependent (Cane et al., 2013; Nicklas and Ward, 1994). Here we reconstituted this behavior *in vitro* using dynamic microtubules and cargo-coupled modules with a defined NDC80 stoichiometry. The tension-dependent modulation of microtubule dynamics at the outer kinetochore by clusters of NDC80 might thus directly contribute to proper microtubule attachment. Stabilization of kinetochore-microtubule interactions by tension has also been recapitulated with purified kinetochore particles from *Saccharomyce cerevisiae* (Akiyoshi et al., 2010). Such endogenous kinetochores, whose homogeneity and stoichiometry of microtubule binders has not been precisely defined, withstand several pN of an externally applied force before detaching from a microtubule (Akiyoshi et al., 2010; Miller et al., 2016; Sarangapani et al., 2014). This, however, appears to be crucially dependent on the Dam1 (DASH) complex, an additional microtubule binder that contributes to the efficient force coupling at the microtubule plus-end in *Saccharomyce cerevisiae* (Lampert et al., 2010; Tien et al., 2010). Dam1, as well as the functionally analogous human Ska complex, also confer plus-end tracking activity *in vitro* to monomeric NDC80 (Janczyk et al., 2017; Lampert et al., 2010; Schmidt et al., 2012; Tien et al., 2010; Welburn et al., 2009). Our observations, however, demonstrate that the constrained stoichiometry and spatial arrangement of multiple NDC80 tethers result in cargo-couplers that follow and modulate microtubule dynamics autonomously. Addressing how additional microtubule binders contribute to enhance the force coupling between kinetochores and dynamic microtubules thus preventing the catastrophic effects of chromosome missegregation on cell physiology is an important task for the future. Here, we precisely constrained a number of NDC80 tethers in a single module to study how multivalency governs the interaction between outer kinetochores and dynamic microtubules. Our work highlights that recapitulating stoichiometry and spatial arrangement of macromolecular assemblies is crucial for the bottom-up reconstitution of biological processes.

## Acknowledgements

We thank I. Stender for assistance with the purification of NDC80 complex, M. Baclayon and R. Dries for assistance with the optical trap setup, N. Taberner for assistance with image analysis and M. Kok and L. Reese for suggestions and discussion regarding the modeling. PJH was supported by an EMBO short-term fellowship (grant 7203). MD acknowledges funding from the European Research Council Synergy Grant MODELCELL (grant 609822) and AM acknowledges funding from the European Research Council Advanced Grant RECEPIANCE (grant 669686) and the Deutsche Forschungsgemeinschaft Collaborative Research Centre (CRC) 1093.

## Author Contributions

All authors designed and interpreted experiments. VAV performed light microscopy, modeling, and optical trapping experiments. PJH prepared and characterized protein complexes and performed electron microscopy. VAV and PJH wrote the manuscript together with MD and AM.

## Material and Methods

### Protein Expression and Purification

Standard Gibson assembly or restriction-ligation dependent cloning techniques were used to generate pLIB vectors with NDC80, NDC80^d80^, NDC80^d^80^ K89A K166A, NUF2, NUF2^K41^A^ K115A, SPC25^SORT^-HIS, SPC24, and SPC24^SPY^. Expression cassettes were combined on pBIG1 vectors using Gibson assembly as described (Weissmann et al., 2016). Baculoviruses were generated in Sf9 insect cells and used for protein expression in Tnao38 insect cells. Between 60 and 72 hours post-infection, cells were washed in PBS (10 mM Na_2_HPO_4_, 1.8 mM KH_2_PO_4_, 2.7 mM KCl, 137 mM NaCl, pH 7.4) and stored at -80 °C. All subsequent steps were performed on ice or at 4 °C. Thawed cells were resuspended in buffer A (50 mM Hepes, pH 8.0, 200 mM NaCl, 5% v/v glycerol, 1 mM TCEP) supplemented with 20 mM imidazole, 0.5 mM PMSF, and protease-inhibitor mix HP Plus (Serva), lysed by sonication and cleared by centrifugation at 108,000g for 30 minutes. The cleared lysate was filtered (0.8 µM) and applied to a 5 ml HisTrap FF (GE Healthcare) equilibrated in buffer A with 20 mM imidazole. The column was washed with approximately 50 column volumes of buffer A with 20 mM imidazole and bound proteins were eluted in buffer A with 300 mM imidazole. Relevant fractions were pooled, diluted 5-fold with buffer A with 25 mM NaCl and applied to a 6 ml ResourceQ column (GE Healthcare) equilibrated in the same buffer. Bound proteins were eluted with a linear gradient from 25 mM to 400 mM NaCl in 30 column volumes. Relevant fractions were concentrated in 30 kDa molecular mass cut-off Amicon concentrators (Millipore) in the presence of additional 200 mM NaCl and applied to a Superose 6 10/300 column (GE Healthcare) equilibrated in 50 mM Hepes, pH 8.0, 250 mM NaCl, 5% v/v glycerol, 1 mM TCEP. Size-exclusion chromatography was performed under isocratic conditions at recommended flow rates and the relevant fraction were pooled, concentrated, flash-frozen in liquid nitrogen, and stored at -80 °C.

Core Traptavidin (T; addgene plasmid #26054) and Dead Streptavidin-SpyCatcher (S; addgene plasmid # 59547) were gifts from Mark Howarth (Chivers et al., 2010; Fairhead et al., 2014) and expressed in E. coli BL21 [DE3] RIPL (Stratagene). Expression was induced in cultures with an OD_600_ of 0.9 by adding 0.5 mM IPTG for 4 hours at 37 °C. Cells were washed in PBS and stored at -80 °C. All subsequent steps were performed on ice or at 4 °C. Cells were thawed in 1,5 volumes of lysis buffer (PBS with 10 mM EDTA, 1 mM PMSF, 1% Triton X-100, 0.1 mg/ml lysozyme), incubated for 60 minutes, lysed by sonication, and centrifuged at 10000*g* for 30 minutes. The supernatant was discarded and pelleted material was resuspended in PBS with 10 mM EDTA and 1% Triton X-100, centrifuged as above, resuspended in PBS with 10 mM EDTA, and centrifuged again. Washed inclusion bodies were resuspended in 6M Guanidine hydrochloride pH 1.0 and cleared at 21130*g* for 10 minutes in 2 ml eppendorf tubes. Protein concentrations were determined using absorbance at 280nm and the denatured T and S subunits were mixed in an approximate 1:2 molar ratio. Refolding into T_x_S_y_ tetramers was accomplished by dropwise dilution and an overnight incubation in a 100x volume of stirring PBS with 10 mM EDTA. This mixture was supplemented with 300 gr (NH_4_)_2_SO_4_ per liter and crudely filtered using paper towels. T_x_S_y_ tetramers were precipitated by doubling the amount of added (NH_4_)_2_SO_4_ and pelleted at 15000*g* or 17000*g* using a JA-14 or JA-10 rotor, respectively. The precipitate was resuspended in 50 mM Boric Acid, 300 mM NaCl, pH 11.0 and dialysed to 20 mM Tris pH 8.0 using a SnakeSkin membrane with a 7 kDa molecular mass cut-off (ThermoFisher). The T_x_S_y_ mixture was loaded onto a 25 ml Source 15Q anion-exchange resin and eluted in 20 mM Tris pH 8.0 with a linear gradient from 100 mM to 600 mM NaCl in 8 column volumes at a flow rate of 1 ml/min. Relevant fractions were pooled, analyzed by SDS-PAGE followed by coomassie staining (boiled and not-boiled), and further purified if required by size-exclusion chromatography using a Superdex 200 10/300 column (GE Healthcare) equilibrated in 20 mM TRIS pH 8.0, and 200 mM NaCl, 2% v/v glycerol, 1 mM TCEP. Purified TS tetramers were concentrated using 10 kDa molecular mass cut-off Amicon concentrators (Millipore), flash-frozen in liquid nitrogen, and stored at -80 °C.

### Assembly of TS-NDC80 modules

A mixture of NDC80 and T_x_S_y_ with a 3-4 fold molar excess of NDC80 per S subunit was incubated for 12-20 hours at 10 °C in the presence of PMSF (1 mM) and protease inhibitor mix (Serva). The formation of T_x_S_y_-SPC24^SPY^_y_ was monitored using SDS-PAGE followed by coomassie staining (samples not boiled). We either used a sortase-labeled fluorescent NDC80 complex or included GGGGK-TMR peptide and a sortase 7M mutant (Hirakawa et al., 2015) in the overnight spy-coupling reaction. In the latter case, molar ratios of approximately 20 and 0.2 compared to NDC80 were used. Reaction mixtures were applied to a Superose 6 increase 10/300 or a Superose 6 increase 5/150 column (GE Healthcare) equilibrated in 20 mM TRIS pH 8.0, 200 mM NaCl, 2% v/v glycerol, 2 mM TCEP. Size-exclusion chromatography was performed at 4 °C under isocratic conditions at recommended flow rates and the relevant fractions were pooled and concentrated using 30 kDa molecular mass cut-off Amicon concentrators (Millipore), flash-frozen in liquid nitrogen, and stored at -80 °C.

### Low-angle metal shadowing and electron microscopy

TS-NDC80 assemblies were diluted 1:1 with spraying buffer (200 mM ammonium acetate and 60% glycerol) and air-sprayed as described (Baschong and Aebi, 2006; Huis In’t Veld et al., 2016) onto freshly cleaved mica pieces of approximately 2x3 mm (V1 quality, Plano GmbH). Specimens were mounted and dried in a MED020 high-vacuum metal coater (Bal-tec). A Platinum layer of approximately 1 nm and a 7 nm Carbon support layer were evaporated subsequently onto the rotating specimen at angles of 6-7° and 45° respectively. Pt/C replicas were released from the mica on water, captured by freshly glow-discharged 400-mesh Pd/Cu grids (Plano GmbH), and visualized using a LaB_6_ equipped JEM-1400 transmission electron microscope (JEOL) operated at 120 kV. Images were recorded at a nominal magnification of 60,000x on a 4k x 4k CCD camera F416 (TVIPS), resulting in 0.18 nm per pixel. Particles were manually selected using EMAN2 (Tang et al., 2007).

### Negative staining and electron microscopy

Taxol-stabilized microtubules were made by polymerizing 20 µM tubulin in the presence of 1 mM GTP at 37°C. Taxol was added to concentrations of 0.2, 2, and 20 µM after 10, 20, and 30 minutes respectively. Stabilized microtubules were sedimented over a warm MRB80 gradient with 40% glycerol, 1 mM DTT, and 20 µM taxol and were resuspended in MRB80 with 20 µM taxol. Microtubules were incubated for 15 minutes at room temperature with 10 nm biotin-nanogold (Cytodiagnostics) and T_1_S_3_[NDC80]_3_. The 10 µL mixture with tubulin at 2 µM, T_1_S_3_[NDC80]_3_ at 0.4 µM, and biotin-nanogold at 0.05 OD (520nm maximum absorbance; 1000x diluted from the stock) was thereafter applied for 45 seconds to freshly glow-discharged 400 mesh copper grids (G2400C, Plano GmbH) with a continuous carbon film. Grids were washed three times with MRB80 buffer, once with freshly prepared 0.75 % uranyl formate (SPI Supplies), and then stained with the uranyl formate for 45 seconds. Excess staining solution was removed by blotting and the specimen was air-dried. Presented micrographs were recorded as described above.

### Tubulin and microtubules

Digoxigenin-labelled tubulin was produced by cycling of porcine brain extract in high-molarity PIPES (Castoldi and Popov, 2003) followed by labelling according to published protocols (Hyman et al., 1991). All other tubulins were purchased from Cytoskeleton Inc. GMPCPP-stabilized seeds were made by two rounds of polymerization to remove any residual GDP. 25 µM tubulin (40% dig-tubulin) supplemented with 1 mM GMPCPP (Jena Biosciences) were polymerized for 30 min at 37°C, spun down in Beckman Airfuge (5 min at 30 psi), resuspended in 75% of the initial volume of MRB80 (80 mM K-Pipes pH 6.9 with 4 mM MgCl_2_ and 1 mM EGTA) and depolymerized on ice for 20 min. After that the solution was supplemented with 1 mM GMPCPP and polymerized again for 30 min at 37°C. Microtubule seeds were sedimented again, resuspended in 50 µL of MRB80 with 10% glycerol, aliquoted and snap-frozen in liquid nitrogen for storage at -80C for up to 2-3 months.

Taxol-stabilized microtubules were made by polymerizing 50 µM tubulin (8% dig-tubulin, 3-6% Hilyte-647 tubulin) in the presence of 1 mM GTP 30 min at 37°C, then 10-25 µM taxol was added for another 30-60 min. Polymerized tubulin was then sedimented in Beckman Airfuge (3 min at 14 psi) and resuspended in 50 µL MRB80 supplemented with 10 µM taxol.

### TIRF microscopy

Coverslips were cleaned in oxygen plasma and silanized as described (Volkov et al., 2014). Flow chambers were constructed by a glass slide, double-sided tape (3M) and silanized coverslip and perfused using a pipet. The chambers were first incubated with ˜0.2 µM anti-DIG antibody (Roche) and passivated with 1% Pluronic F-127. Then taxol-stabilized microtubules (300 µL diluted 1:30-1:600) were introduced followed by the reaction mix and the chambers were sealed with valap. The reaction mix contained MRB80 buffer with 1 mg/ml-casein, 10 µM taxol, 4 mM DTT, 0.2 mg/ml catalase, 0.4 mg/ml glucose oxidase and 20 mM glucose, supplemented with 10-35 pM of TS-NDC80. For experiments with dynamic microtubules the taxol microtubules were substituted with GMPCPP-stabilized microtubule seeds, and the reaction mix (short of taxol) was supplemented with 8 µM tubulin (4-6% labeled with HiLyte-647), 1 mM GTP and 0.1% methylcellulose. In all experiments the reaction mix was centrifuged in Beckman Airfuge for 5 min at 30 psi before adding to the chamber.

Imaging was performed at 30°C using Nikon Ti-E microscope (Nikon) with the perfect focus system (Nikon) equipped with a Plan Apo 100X 1.45NA TIRF oil-immersion objective (Nikon), iLas^2^ ring TIRF module (Roper Scientific) and a Evolve 512 EMCCD camera (Roper Scientific). Images were acquired with MetaMorph 7.8 software (Molecular Devices). The final resolution was 0.16 µm/pixel. The objective was heated to 34°C by a custom-made collar coupled with a thermostat, resulting in the flow chamber being heated to 30°C.

### Image analysis

All images were analysed using Fiji (Schindelin et al., 2012). Kymographs were produced by a custom macro that creates an average projection perpendicular to a selected line through a reslice operation. Resulting kymographs were then analysed using a custom script in MatLab R2013b. Each horizontal line of the kymograph was fitted with a Gaussian function, with its peak being the central position of the fluorescent spot, and the area under the curve being the spot intensity. Fluorescence intensity of the spot before the first bleach event was averaged to obtain the initial intensity of the spot. Height of the individual bleach event was determined by obtaining the bleaching traces of TS-NDC80 modules diffusing on taxol-stabilized microtubules, then smoothing these traces with the Chung-Kennedy filter as described (Chung and Kennedy, 1991; Reuel et al., 2012). Resulting smoothed traces were then used to build the histograms of all intensities that occurred during bleaching and further analysed as described (Grishchuk et al., 2008). Lifetime measurements were performed by calculating the time difference between the landing and detaching events in kymographs. For chambers containing trivalent and tetravalent TS-[NDC80] modules, in which the oligomers were bound to the microtubules immediately after addition to the chamber, the amount of time between addition of the oligomer and start of imaging (1-3 min) was neglected, and spots already present on the microtubules in the first frame were considered as just landed.

### Preparation of beads coated with TS-NDC80 modules

1 µm glass COOH-functionalized beads (Bangs Laboratories) were suspended by sonication as 1% w/v in MES buffer (25 mM MES pH 5 supplemented with 0.05% tween-20), washed by centrifugation at 16 kG for 1 min, and then activated with EDC and Sulfo-NHS, each at 10 mg/ml, for 30 min at 23°C with vortexing. After 3 washes the beads were allowed to bind a mixture of 2 mg/ml PLL-PEG (Poly-L-lysine (20 kDa) grafted with polyethyleneglycole (2 kDa), SuSoS AG) with 0-10% v/v of PLL-PEG-biotin for 30 min at 23C. The reaction was quenched by adding 200 mM glycine. The beads were washed 3 times, resuspended at 0.2% w/v and stored at 4C.

Before each experiment, 10 µL of PLL-PEG-coated beads were washed using washing buffer (MRB80 with 2 mM DTT and 0.4 mg/ml casein), resuspended in 10 µL and mixed with 10 µL of NDC80C oligomer in the same buffer. Incubation was performed for 1 hr on ice with frequent pipetting, then the beads were washed 3 times and resuspended in 30 µL of washing buffer. Flow chambers with GMPCPP-stabilized seeds were prepared as described above, the reaction mix contained MRB80 buffer with 10-12 µM tubulin, 1 mM GTP, 1 mg/ml-casein, 4 mM DTT, 0.2 mg/ml catalase, 0.4 mg/ml glucose oxidase and 20 mM glucose. This reaction mix was centrifuged in Beckman Airfuge for 5 min at 30 psi, and then 1 µL of beads suspension was added to 14 µL of the reaction mix, added to the chamber and the chamber was sealed with valap. The final concentration of the beads was 0.004% w/v (80 fM, without losses).To measure bead brightness, the beads were washed twice in MRB80 buffer, resuspended in 10 µL and a 4 µL drop was placed on a plasma-cleaned coverslip. The coverslip was then put on top of the slide and sealed with valap. Beads were imaged using brightfield and laser epifluorescence to get the positions and fluorescence intensities of all beads, respectively. Fluorescence intensity of a single fluorophore for normalization of bead fluorescence was obtained as described (Volkov et al., 2014).

### Laser tweezers and experiments with the beads

DIC microscopy and laser tweezers experiments were performed using a custom-built instrument described elsewhere (Baclayon et al., 2017). Images were captured using Andor Luca R or QImaging Retiga Electro CCD cameras and MicroManager 1.4 software. At the start of each experiment 50 frames of bead-free fields of view in the chamber were captured, averaged, and used later for on-the-fly background correction. The images were acquired at 8 frames per second, subjected to background substraction and each 10 consecutive frames were averaged. Calibration of quadrant photo-detector (QPD) response was performed by sweeping the trapping 1064 nm beam with the trapped bead across the tracking 633 nm beam with acousto-optic deflector (AOD) over the distance of ± 400 nm in two orthogonal directions. The central ± 200 nm region of the resulting voltage-displacement curve was then fitted with a linear fit to determine the conversion factor. Stiffness calibration was performed by fitting the power spectral density as described (Tolic-Norrelykke et al., 2004) with correction for the proximity of the coverslip (Nicholas et al., 2014). The axial position of the free bead in a trap was adjusted to leave 100-200 nm between the surfaces of the bead and the coverslip, while having the coverslip-associated microtubules in the focal plane of the objective. QPD signal was sampled at 100 or 10 kHz without additional filtering. All experiments were performed at 0.2-0.4W of the 1064 nm laser resulting in a typical trap stiffness of 0.015-0.033 pN/nm.

Force signals were analysed in MatLab R2013b. Direction of the force development was checked for consistency with video recording for each signal. Signals with ambiguous direction of the force, or beads attached to more than one microtubule were discarded. X and Y coordinates from the QPD recordings were then rotated to correspond to the directions along and across the microtubule. A portion of the signal corresponding to microtubule pulling was downsampled to 1 kHz and smoothed with a Chung-Kennedy filter (Chung and Kennedy, 1991; Reuel et al., 2012). The stall force was determined as the difference between the stall level and the free bead level in the smoothed signal along the microtubule multiplied by trap stiffness, corrected for the nonlinear increase of the force as a function of the distance from the trap center (Simmons et al., 1996).

To calculate the effective stiffness of the link between the bead and the microtubule, we have measured the variance <var> of the signal along the microtubule during stall (for microtubules pulling on the bead), or during a pause in the piezo-stage motion (for control experiments). Stiffness was then calculated as *k*_*B*_*T/<var>*.

### Computer simulation of lifetimes

Simulations were performed using Gillespie algorithm (Gillespie, 1977) in MatLab R2013b. Each simulation run started with an oligomer with N binding sites with only one binding site being attached. In the case of N = 1 the only event that can occur afterwards is detachment with a fixed rate *k*_*off*_ determined as 1/(average lifetime of T_3_S_1_x[NDC80C]_1_) = 0.62 s^−1^. If N > 1, detachment of each “attached” binding site happens at a rate *k*_*off*_, and attachment of each available site inside the oligomer happens at a rate *k*_*on*_. Simulation stopped when all binding sites in an oligomer transitioned to the “detached” state. The time elapsed until the next event was determined by calculating the propensity towards this event using pre-generated random numbers.

## Figure legends

**Figure S1.**
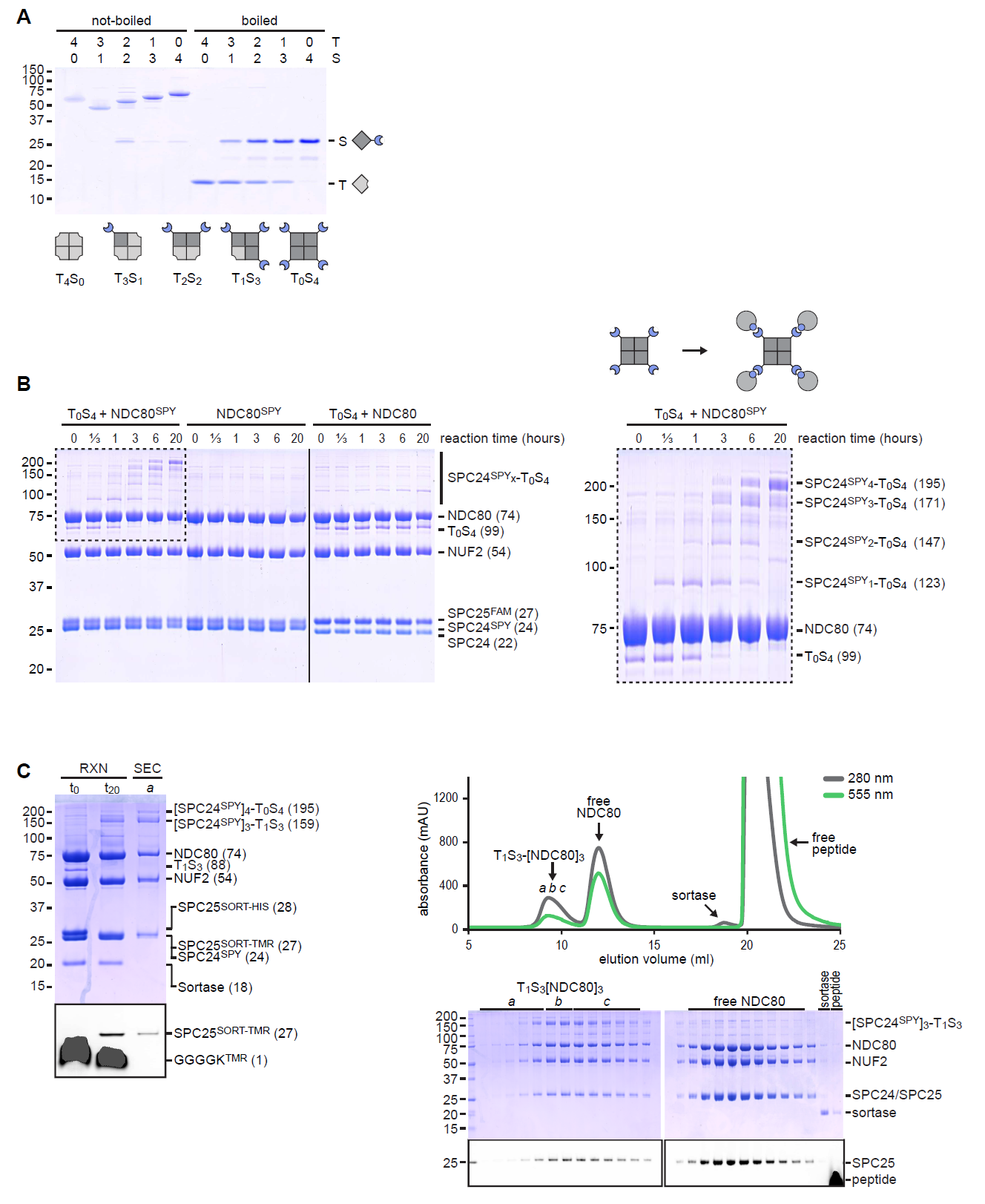
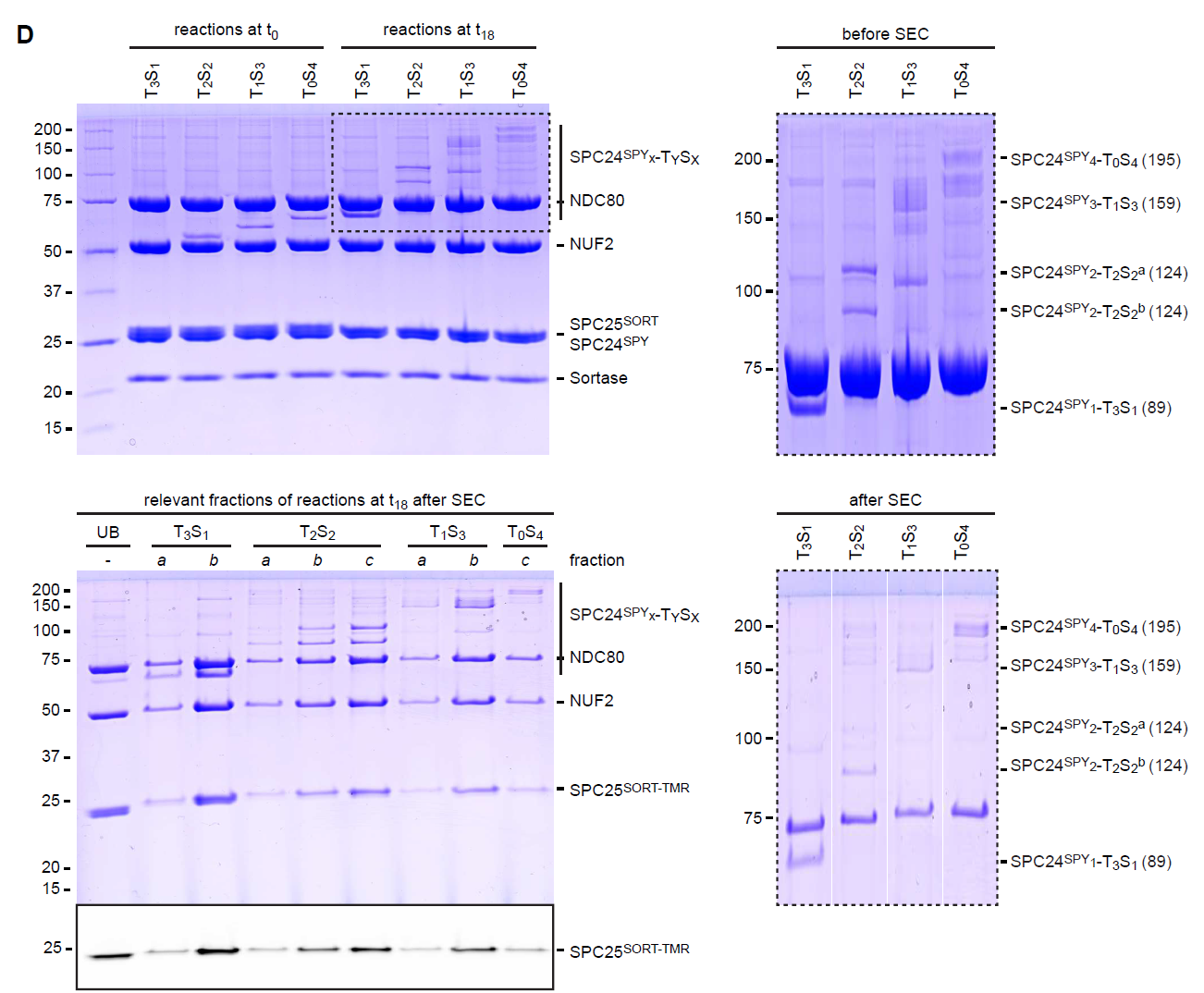
A reconstituted system to precisely control NDC80 stoichiometry. **a**) T_y_S_x_ variants were separated by ion-exchange chromatography based on the pI difference of T (5.1) and S (4.5). Collected assemblies were analyzed by SDS-PAGE as tetramers (not-boiled) or in a denatured form (boiled). **b**) SDS-PAGE analysis (samples not-boiled) to monitor the formation of S-SPC24^SPY^ complexes. NDC80 complexes without added T_0_S_4_ or without a SPY-tag were analysed as a control. The boxed area is shown at larger magnification on the right. **c**) Samples before (t_0_) and after (t_20_) the reaction were analysed by SDS-PAGE (samples not-boiled) to monitor coupling of SPC24^SPY^ to T_1_S_3_ tetramers and fluorescent labelling of SPC25^SORT^. Size exclusion chromatography was used to separate T_1_S_3_[NDC80]_3_ assemblies from sortase and from the excess of unreacted NDC80 and free peptide. The lower panels show the GGGGK^TMR^ and SPC25^TMR^ in-gel fluorescence. **d**) Samples before (t_0_) and after (t_18_) the reaction were analysed by SDS-PAGE (samples not-boiled) to monitor coupling of SPC24^S^PY to T_y_S_x_ tetramers and fluorescent labelling of SPC25^SORT^. Size-exclusion chromatography was used to separate T_y_S_x_[NDC80]_x_ assemblies from sortase and from the excess of unreacted NDC80 and free peptide. Fractions *a*, *b*, and *c*, indicate the front, middle, and tail of the peak as indicated in Figure S1C.

**Figure S2.**
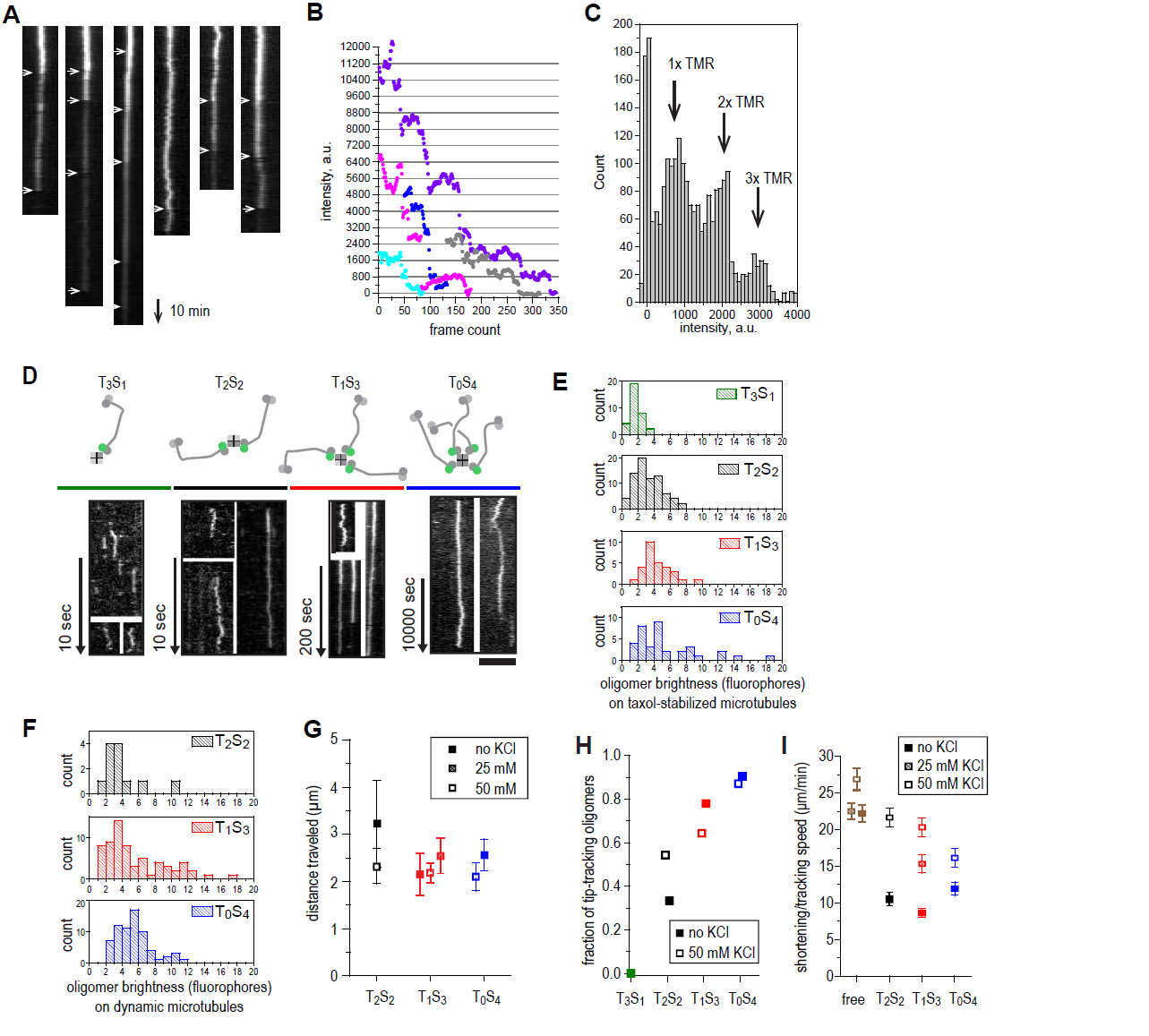
Characterization of TS-NDC80 assemblies on taxol-stabilized and dynamic microtubules. **a**) Kymographs showing individual bleaching traces of T_0_S_4_-[NDC80^TMR^]_4_ oligomers attached to taxol-stabilized microtubules. Arrows show bleaching events See Materials and Methods for description. **b**) Fluorescence intensities of several photobleaching curves. **c**) Histogram of all intensities that occurred during bleaching of 13 microtubule-bound T_0_S_4-_[NDC80^TMR^]_4_ oligomers. Arrows show positions of individual peaks corresponding to 1, 2 and 3 TMR fluorophores. **d**) Kymographs of mono-, di-, tri-, and tetravalent NDC80 complexes binding to taxol stabilized microtubules. Scale bar 5 µm. **d**) Brightness distribution of TS-NDC80 assemblies on taxol-stabilized microtubules. **e**) Brightness distribution of TS-NDC80 assemblies on dynamic microtubules. **f**) Distance travelled by TS-NDC80 modules moving with the tips of the shortening microtubules. **g-h**) Presence of 25-50 mM KCl in the motility buffer does not prevent TS-NDC80 assemblies from tip-tracking and slowing down the microtubule shortening. Data are shown as mean ± SEM.

**Figure S3.**
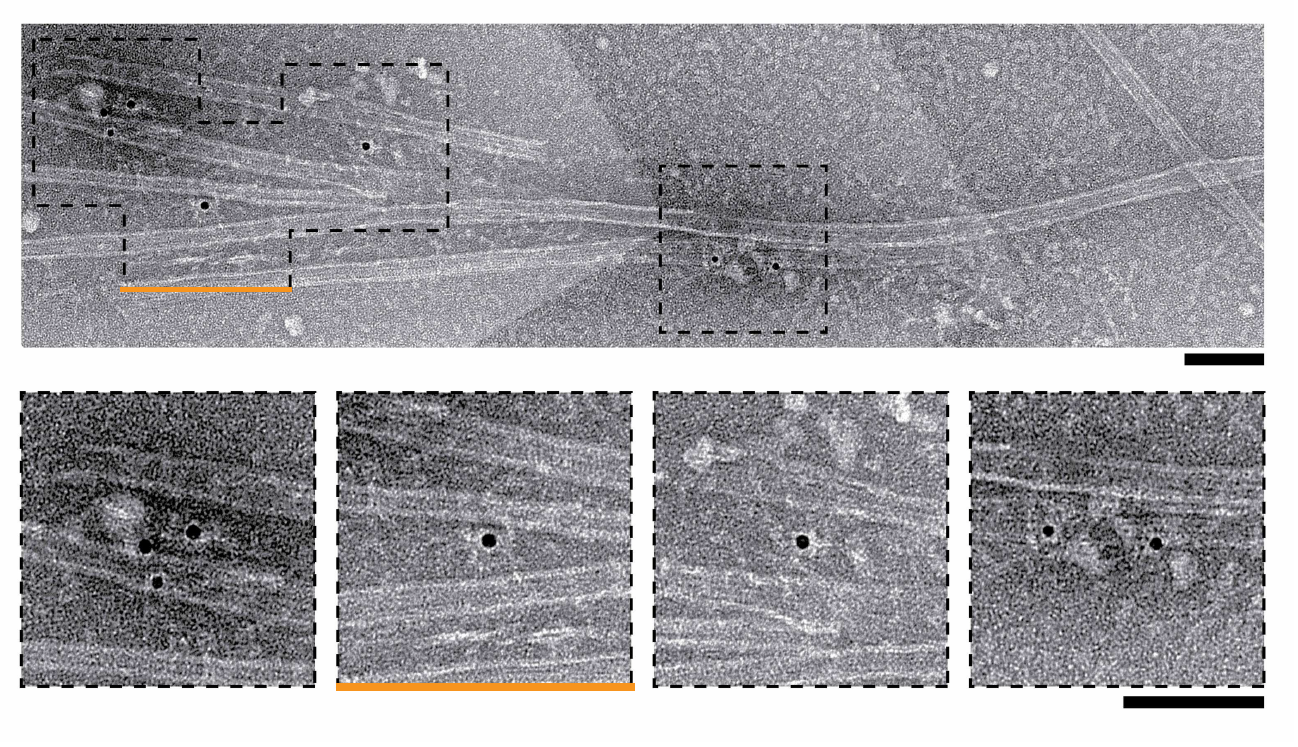
Negative-stain EM of microtubules and nanogold particles coated with T_1_S_3_[NDC80]_3_. Boxed areas in the upper micrograph are shown below at a higher magnification. The orange line marks the micrograph shown in main Figure 2E. Scale bars 100 nm.

**Figure S4.**
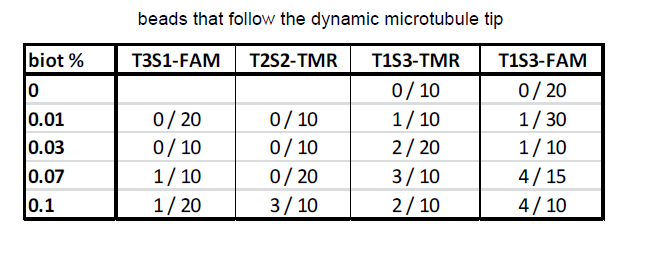
Tip-tracking events for differently coated beads. Accompanying Figure 2I.

## Video legends

**Video 1 Single-molecule TIRF microscopy of TS-NDC80 modules on dynamic microtubules**. 35 pM of T_0_S_4_[NDC80^TMR^]_4_ (green) in the presence of 8 µM tubulin labelled with HiLyte-642 (red) and in the absence of KCl. The two-channel images were acquired every 1.1 s (shown at 30 fps). Top left corner shows time in min:sec. White arrows mark tip-tracking events. Scale bar 5 µm.

**Video 2 A disassembling microtubule tip pulls on a trapped bead**. A bead coated with 3% PLL-PEG-biotin and then saturated with T_2_S_2_[NDC80^TMR^]_2_ was attached to a microtubule with a trap (see also Figure 3B for still images and a complete QPD trace of this signal). Images were acquired at 8 fps in DIC, background subtracted, and each 10 consecutive frames averaged (see Materials and Methods). The timer at the bottom left corner shows elapsed time in seconds. The microtubule continues growing until about 230s, when it switches to shortening and pulls on the bead at 248s (evident from the bead’s brief displacement in the direction of microtubule disassembly). From 265s till 290s the microtubule end is seen disassembling to the left of the bead. Scale bar 5 µm.

**Video 3 A microtubule is rescued five times at the bead attachment site**. A bead coated with 0.3% PLL-PEG-biotin and then saturated with T_1_S_3_[NDC80^FAM^]_3_ was attached to a microtubule with a trap. The microtubule experiences dynamic instability, but its shortening is five times in a row rescued at the attached bead (at 2, 9, 24, 47 and 60 min from the start of the experiment, see timer in the top left corner). Note that each rescue is preceded by a displacement of the bead along the microtubule axis in the direction of disassembly. After 63 minutes, the trap stiffness is increased from initial 0.03 pN/nm to 0.13 pN/nm to manually remove the bead from the microtubule. The microtubule without the attached bead depolymerizes without stalling or rescue events. Scale bar 5 µm.

